# CRaTER enrichment for on-target gene-editing enables generation of variant libraries in hiPSCs

**DOI:** 10.1101/2023.01.25.525582

**Authors:** Clayton E. Friedman, Shawn Fayer, Sriram Pendyala, Wei-Ming Chien, Linda Tran, Leslie Chao, Ashley Mckinstry, Elaheh Karbassi, Aidan M. Fenix, Alexander Loiben, Charles E. Murry, Lea M. Starita, Douglas M. Fowler, Kai-Chun Yang

## Abstract

Standard transgenic cell line generation requires screening 100-1000s of colonies to isolate correctly edited cells. We describe <u>CR</u>ISPR<u>a</u> On-<u>T</u>arget <u>E</u>diting <u>R</u>etrieval (CRaTER) which enriches for cells with on-target knock-in of a cDNA-fluorescent reporter transgene by transient activation of the targeted locus followed by flow sorting to recover edited cells. We show CRaTER recovers rare cells with heterozygous, biallelic-editing of the transcriptionally-inactive *MYH7* locus in human induced pluripotent stem cells (hiPSCs), enriching on average 25-fold compared to standard antibiotic selection. We leveraged CRaTER to enrich for heterozygous knock-in of a library of single nucleotide variants (SNVs) in *MYH7*, a gene in which missense mutations cause cardiomyopathies, and recovered hiPSCs with 113 different *MYH7* SNVs. We differentiated these hiPSCs to cardiomyocytes and show MYH7 fusion proteins can localize as expected. Thus, CRaTER substantially reduces screening required for isolation of gene-edited cells, enabling generation of transgenic cell lines at unprecedented scale.

## INTRODUCTION

Gene editing with programmable endonucleases occurs via two endogenous repair mechanisms following the generation of a DNA double-strand break (DSB). The dominant DNA repair pathway, non-homologous end joining (NHEJ), is error-prone and typically results in small nucleotide insertions or deletions at the DSB (1), errors that can be leveraged to knock out genes of interest (GOI) by shifting the open reading frame to introduce a premature stop codon. Alternatively, homology-directed repair (HDR) recombines homologous DNA at the DSB and can be leveraged to knock-in exogenous transgenic single-stranded oligodeoxyribonucleic (ssODN) or plasmid DNA donor repair templates. HDR rates are low in mammalian cells and substantially lower in pluripotent stem cells (2) despite extensive efforts to boost HDR efficiency (3,4). Moreover, low transcriptional activity of recombined transgenes within transcriptionally-inactive genes or ‘safe harbor’ loci (5,6) can reduce constitutive expression of selectable markers, such as genes encoding fluorescent proteins or conferring antibiotic resistance, resulting in incomplete selection that requires additional colony screening to isolate bona fide on-target gene-edited cells (7). This laborious step in turn limits the scale of transgenic cell line generation and the feasibility of more complicated genetic modifications, such as biallelic editing or variant library generation.

In addition to augmenting HDR rates, various complementary approaches have emerged in the last decade to facilitate enrichment of on-target gene editing. However, many of these approaches rely on using a previously generated locus-specific ‘landing pad’ cell line prior to introducing desired transgenes, limiting widespread utility (8–10). Reporters of transfection, such as co-targeting with selection (11) or transient reporters of editing efficiency (12) improve selection of cells transfected with the required gene-editing machinery (and thus cells more likely to be gene-edited), yet these methods do not elicit direct readouts of genetic modification to the targeted GOI and predominately enrich for cells with homozygous gene-edits. A method described by Roberts *et al*. enables the generation of human induced pluripotent cell (hiPSC) lines with fluorescently tagged transcriptionally inactive genes (13), however, this approach relies on robust expression of an intermediate fluorescent reporter for enrichment. Another method developed by Arias-Fuenzalida *et al*. enriches for cells using positive and negative selection modules in the repair template while introducing single nucleotide variants (SNV) via the homology arm (14), yet this approach was only tested for transcriptionally-active genes in hiPSCs; the rate of gene-editing in transcriptionally-inactive genes using this approach remains unknown. Ultimately, these enrichment strategies rely on robust expression of a constitutive fluorescent reporter which will vary depending on native transcriptional activity of the targeted genetic locus (5,6).

Here, we report a novel method called <u>CR</u>ISPR<u>a</u> On-<u>T</u>arget <u>E</u>diting <u>R</u>etrieval (CRaTER) to enrich for on-target gene-edited cells irrespective of transcriptional activity at the targeted GOI. We chose a low probability gene-editing scenario to examine the enrichment capabilities of CRaTER: Heterozygous, biallelic-editing of the transcriptionally inactive *MYH7* locus in hiPSCs. *MYH7* encodes myosin heavy chain β (MHC-β), a sarcomeric thick filament protein that is not expressed in hiPSCs (15). We use CRISPR/Cas9 to knock-in transgenes containing a promoterless partial *MYH7* cDNA (exon 22-40) fused with a C-terminal fluorescent protein gene (eGFP or mTagBFP2) in *cis* with an antibiotic resistance cassette (PGK-PuroR or EF1α-BSD) into the endogenous *MYH7* locus. Following standard antibiotics selection, we transiently overexpress *MYH7* in hiPSCs using CRISPRa of the endogenous *MYH7* promoter and sort for eGFP^+^/mTagBFP2^+^ cells using fluorescence-activated cell sorting (FACS), enabling 9-fold enrichment of desired heterozygous, biallelic editing.

Heterozygous, pathogenic missense mutations in *MYH7* cause autosomal dominant hypertrophic and dilated cardiomyopathy (16,17) and also skeletal myopathies (18). To enable future high-throughput variant effect studies of *MYH7* missense variants, we generated a library of heterozygous *MYH7* SNVs in hiPSCs by mutagenizing five amino acid positions to saturation and leveraged CRaTER to enrich for hiPSCs with 78 heterozygous, biallelic missense variants in the endogenous *MYH7* locus – a 38.6-fold enrichment compared to antibiotics selection alone, yielding a library of cells with 90% on-target gene-edits. hiPSCs enriched using CRaTER maintain pluripotency, demonstrate proper localization of MHC-β fusion proteins, and differentiate to functional cardiomyocytes. Together, CRaTER enrichment reduces the amount of colony screening needed by >10-fold, enabling the generation of substantially more correctly gene-edited hiPSCs lines compared to contemporary methods.

## MATERIAL AND METHODS

### Cell culture and maintenance

All cells maintained at 37° C in 5% CO_2_. Human induced pluripotent stem cells (hiPSC), including wildtype (WT) WTC11 hiPSCs (XY; a gift from Bruce Conklin) and subclones derived from these cells were cultured on Matrigel (Corning; cat. no. 354277) and fed every other day with stem cell media: mTeSR Plus with supplement (STEMCELL Technologies; cat. no. 100-0276) and 0.5% penicillin-streptomycin (ThermoFisher; cat. no. 15140122). When 80% confluent (approximately every 4 days) hiPSCs were dissociated using 0.5 mM EDTA (Invitrogen; cat. no. 15575-038) in DPBS (Invitrogen; cat. no. 14190250) and resuspended in stem cell media supplemented with 10 μM Y-27632 dihydrochloride (Rho kinase [ROCK] inhibitor; Tocris; cat. no. 1254). hiPSCs were used for no more than 10 passages after thawing. hiPSC subclones were isolated following sparse plating (~88 cells/cm^2^) by manually picking colonies and further expanded for genotyping and/or maintenance.

### Cardiac-directed differentiation

Small molecule cardiac-directed differentiation was performed following previous protocols (15) with adaptions. On differentiation day T-1, hiPSCs were dissociated as above and replated into a 24 well-plate at 7.89 × 10^5^ cells/cm^2^. On T0 (activation day), the stem cell media with ROCK inhibitor was removed, cells washed once with DPBS, and fed with RBA media: RPMI (Invitrogen; cat. no. 11875135), 0.5 mg/ml bovine serum albumin (Sigma; cat. no. A9418), 0.213 mg/ml ascorbic acid (Sigma; cat. no. A8960), and 0.5% penicillin-streptomycin supplemented with 4 μM CHIR-99021 (Cayman; cat. no. 13122). On T3, CHIR-99021-containing media was removed, cells washed once with DPBS, and fed with RBA supplemented with 2 μM Wnt-C59 (Selleck; cat. no. S7037). On T5, Wnt-C59-containing media was replaced with RBA. On T7 (and every other day afterwards), media was replaced with cardiomyocyte media: RPMI, B27 plus insulin (Invitrogen; cat. no. 17504044), and 0.5% penicillin-streptomycin.

To enrich for cardiomyocytes using lactate selection, T19-21 cells were washed once with DPBS and then dissociated using 37° C 0.5 mM EDTA with 0.5% trypsin (Invitrogen; cat. no.15090046) in DPBS for 7 minutes at 37° C in 5% CO_2_. The cells were collected, 25 μU DNAse I (Sigma; cat. no. 260913) was added, and then cells were triturated using a serological pipette until homogenized. The reaction was stopped using an equal volume of cardiomyocyte media supplemented with 5% fetal bovine serum (FBS; Biowest: cat. no. S1520) and filtered using a 100 μm strainer. Cells were centrifuged at 300 × *g* for 3 minutes and the supernatant aspirated then the pellet was resuspended in cardiomyocyte media with 5% FBS and replated at 5.3-8.8 × 10^5^ cells/cm^2^ in 10 cm plates and incubated overnight at 37° C in 5% CO_2_. For the next 4 days, cells were fed daily with lactate media: DMEM (Invitrogen; cat. no. A1443001), 2 mM L-glutamine (Invitrogen; cat. no. 25030081), 0.5% penicillin-streptomycin, and 4 mM sodium L-lactate (Sigma; cat. no. 71718).

### Plasmid DNA repair template cloning

All plasmids purified with phenol/chloroform before concentration quantification and transfection.

#### Cloning of the pJet-MYH7-eGFP-PGK-PuroR repair template vector

The 5’ homology arm including *MYH7* intron 21 with a 7-nucleotide deletion (GGAGGGG) in the sgRNA sequence to prevent re-cutting and the 3’ homology arm were isolated by PCR using WT WTC11 gDNA as template. The 5’ and 3’ homology arms were extended 500 bp upstream and downstream of *MYH7* targeting sgRNA sites, respectively. The 3’ *MYH7*-eGFP and SV40pA fragments were isolated from a full-length *MYH7* cDNA cloned in-frame into pEGFP-N1 (Clontech Laboratories) by PCR with a SalI site at 3’-end. The 5’ homology arm-intron 21 and the 3’ cDNA *MYH7*-eGFP-SV40pA was joined by PCR and cloned into pJET1.2 (ThermoFisher, cat. no. K1231). The mPGK-PuroR-bGHpA fragment was isolated from p2attPC (Addgene 51547) with the mCherry deleted by PCR. The mPGK-Puro-bGHpA and 3’ homology arm were joined by PCR with primers containing a SalI site at 5’-end of mPGK and cloned into pJET1.2. Finally, the 5’ homology arm and 3’ cDNA-eGFP-SV40pA was isolated and cloned into the mPGK-Puro-bGHpA-3’ homology arm at NotI and SalI sites.

#### Cloning of the pJet-MYH7-mTagBFP2-EF1α-BSD repair template vector

The pJet-MYH7-eGFP-PGK-PuroR vector was used as template to generate the pJet-MYH7-mTagBFP2-EF1α-BSD vector. Briefly, inverse PCR was used to first delete PGK-PuroR and create a linearized vector with terminal ends sharing homology with EF1α-BSD amplified from lentiCas9-Blast vector (Addgene 52962). *In vivo* assembly (IVA) (19) was used to ligate these fragments and create an intermediate vector, pJet-MYH7-eGFP-EF1α-BSD. This vector was then used as template in inverse PCR to delete eGFP and replace with mTagBFP2 amplified from a custom synthesized vector. IVA was used to ligate these fragments and generate the pJet-MYH7-mTagBFP2-EF1α-BSD plasmid which was verified using Sanger sequencing.

### Mutagenesis of the pJet-MYH7-mTagBFP2-EF1α-BSD vector

The library was cloned using IVA (19) and inverse PCR with degenerate primers as previously described (20). Briefly, a forward primer containing a 5’ homology region, a degenerate NNK codon at the targeted amino acid position, and a 3’ extension region was designed for each of the *MYH7* exon 22 positions targeted: 848, 850, 852, 865, and 866. Reverse primers were reverse complements of the 5’ homology region. A separate PCR reaction was performed for each position using the pJet-MYH7-mTagBFP2-EF1α-BSD plasmid backbone as template with Q5 polymerase (NEB; cat. no. M0491S) and 5 μM bespoke forward and reverse primers (IDT), and then digested overnight with DpnI at 37° C (NEB; cat. no. R0176S). 3 μL of digested linear PCR product was transformed into NEB stable competent *E. coli* (NEB; cat. no. C3040H) and grown overnight in 3 mL LB-ampicillin (100 μg/ml), followed by separate midiprep (Sigma; cat. no. NA0200-1KT). Specific single nucleotide variant (SNV) plasmid repair templates were generated in the same manner, but using a single set of bespoke inverse primers for IVA (see **Table 1**).

**Table 1.**
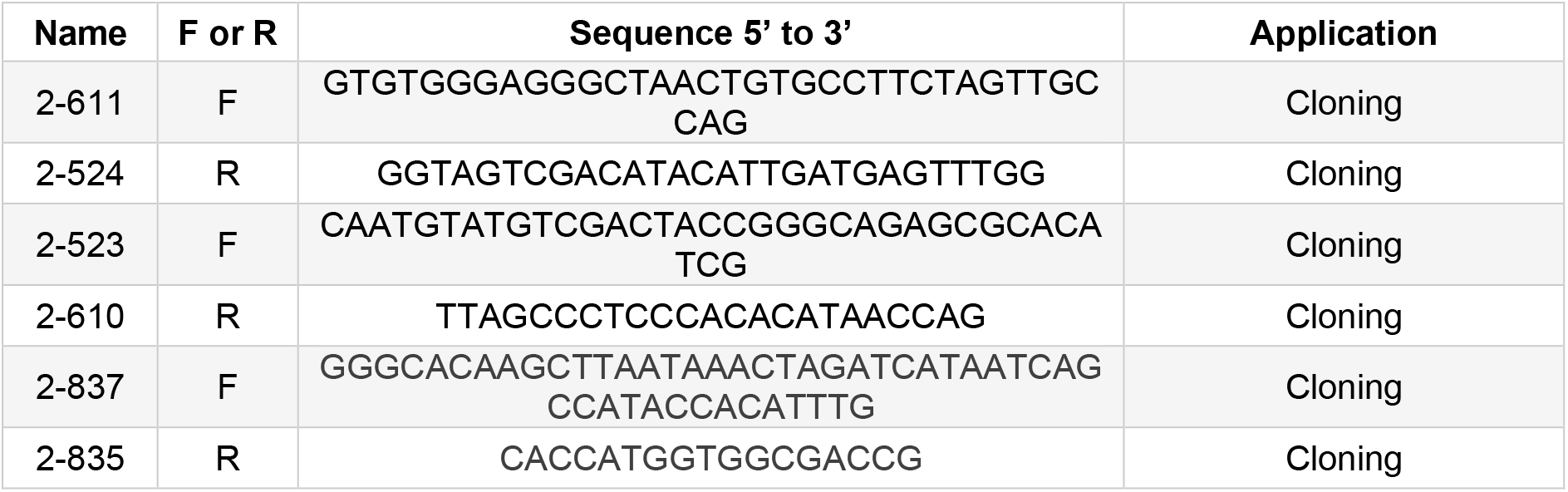
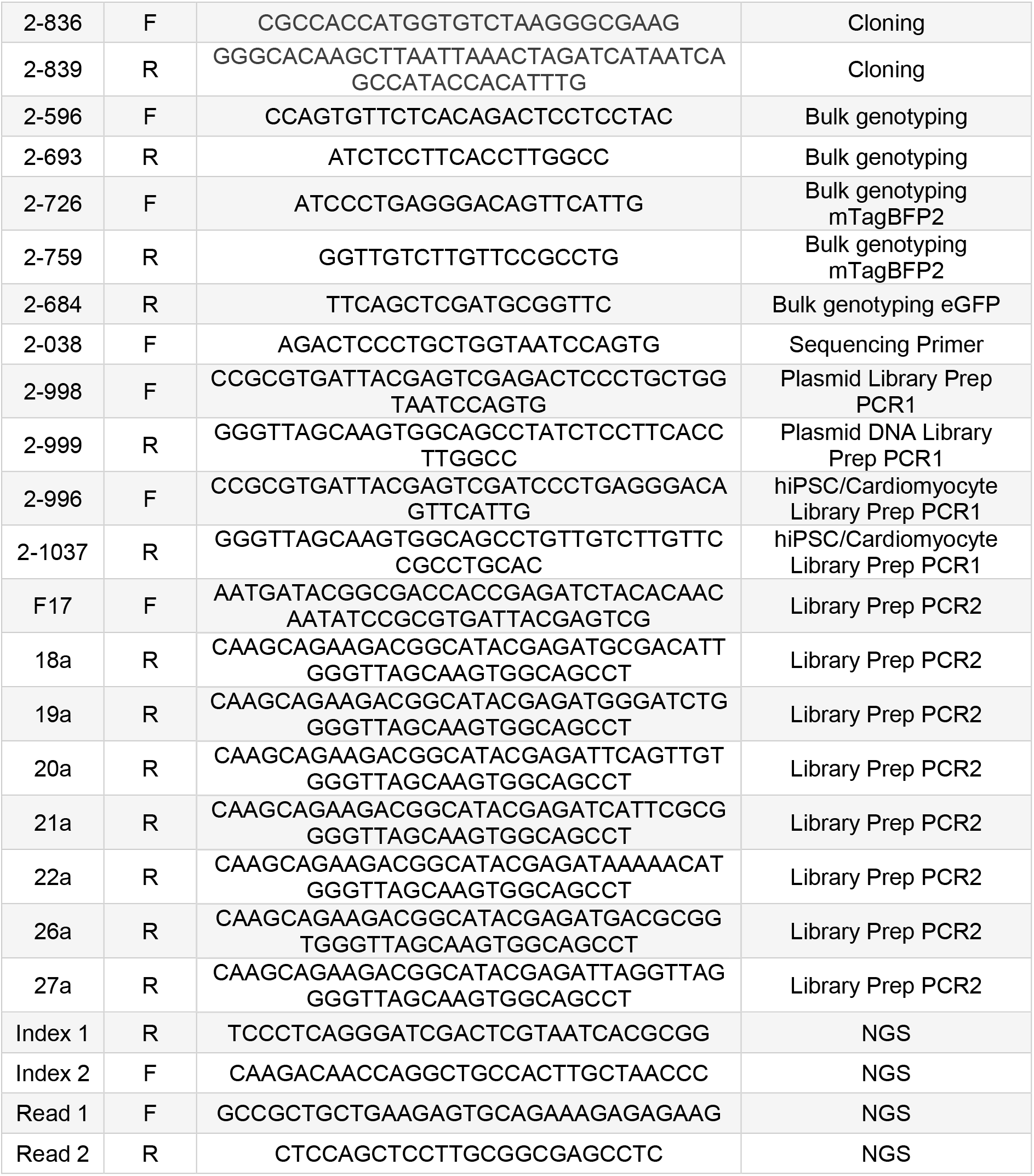
List of primers used in this study.

### Synthetic sgRNA design for CRISPR/Cas9 and CRISPRa

To design synthetic sgRNA for CRISPR/Cas9 gene-editing, we input 150 bp DNA sequences of *MYH7* intron 21 and 24 separately into CCTop and selected sgRNA with high CRISPRater scores and minimal predicted off-targets (21,22). To design synthetic sgRNA for CRISPRa activation of *MYH7* during CRaTER, we input 150 bp DNA sequences upstream of the transcription start sites of *MYH7* into CCTop and followed similar selection criteria as above.

### hiPSC gene editing

#### Generation of CLYBL^dCas9-VPR/dCas9-VPR^ hiPSCs

CLYBL^dCas9-VPR/dCas9-VPR^ cells were generated using the WT WTC11 hiPSC line (a gift from B. Conklin), targeting the endogenous *CLYBL* locus between exons 2 and 3. DNA plasmids used for *CLYBL* locus targeting were pC13N-iCAG.copGFP (Addgene 66578), pZT-C13-L1 (Addgene 62196), pZT-C13-R1 (Addgene 62197) (23). DNA plasmids used for transgene cloning were lenti-EF1α-dCas9-VPR-Puro (Addgene 99373) (24) and IGI-P0492 pHR-dCas9-NLS-VPR-mCherry (Addgene 102245). To generate plasmid DNA repair template for expression of dCas9-VPR driven by a constitutive CAG promoter, dCas9-VPR-P2A-mCherry was cloned into the pC13N-iCAG.copGFP backbone, replacing the copGFP sequence. Plasmid sequences were verified by Sanger sequencing. WT WTC11 hiPSCs were co-transfected with the pC13N-iCAG-dCas9-VPR-P2A-mCherry repair template and pZT-C13-L1 and pZT-C13-R1 plasmids using GeneJuice transfection reagent (Sigma; cat. no. 70967). Edited hiPSCs were selected using G418 (InvivoGen; cat. no. ant-gn-1). Two weeks after transfection, individual clones were isolated and genotyped. The clone used in this study has homozygous knock-in of the transgene at the *CLYBL* locus without random integration of the plasmid detected by PCR.

#### Generation of individual, heterozygous *MYH7* SNV hiPSC lines

CRISPR/Cas9 gene-editing and electroporation were used to biallelically edit the endogenous *MYH7* locus in the CLYBL^dCas9-VPR/dCas9-VPR^ WTC11 hiPSC genetic background (see above). Briefly, we used the Neon Transfection System (ThermoFisher; cat. no. MPK5000) with optimized parameters for the 10 μL tip (1400 V, 20 ms, 1 pulse) to transfect 5 × 10^5^ CLYBL^dCas9-VPR/dCas9-VPR^ hiPSCs with 1 μL of each 30 nM synthetic sgRNA (Synthego; see **Table 2**) targeting *MYH7* intron 21 and 24, 1 μL of 20 μM *Sp*Cas9 2NLS protein (Synthego), 1.5 μg of pJet-MYH7-eGFP-PGK-PuroR (WT *MYH7*) plasmid DNA, and 1.5 μg of pJet-MYH7-mTagBFP2-EF1α-BSD (variant *MYH7*) plasmid DNA, in addition to 300 ng of pEF1α-BCL-XL plasmid DNA to promote survival (25). Following transfection, cells were plated into stem cell media without penicillin-streptomycin supplemented with CloneR2 (STEMCELL Technologies; cat. no. 100-0691) and 10 μM ROCK inhibitor. Media was replaced the following day and cells allowed to recover for an additional 48 hours. Cells were then expanded and a cell pellet was harvested for pre-antibiotic bulk genotyping. The remainder of cells were fed daily for 48 hours with stem cell media containing 1.25 μg/mL blasticidin S HCl (ThermoFisher; cat. no. BP264725) and 0.175 μg/mL puromycin dihydrochloride (ThermoFisher; cat. no. A1113803). Cells were then expanded and a cell pellet was harvested for post-antibiotic bulk genotyping, split at 88 cells/cm^2^ into a 10 cm plate for colony genotyping, and expanded for CRaTER enrichment of edited cells.

**Table 2.**
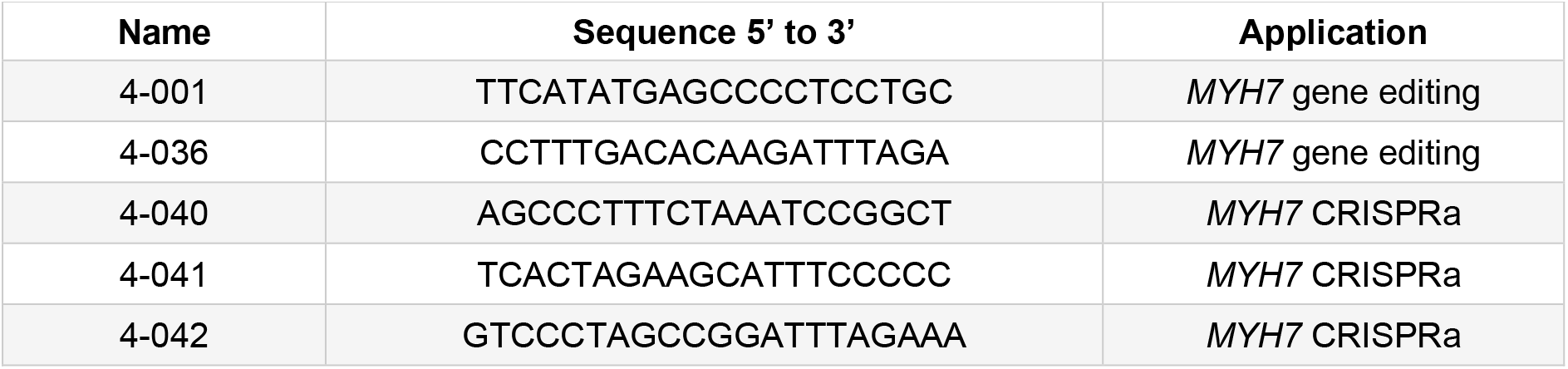
List of synthetic sgRNA used in this study.

#### Pooled generation of a library of single, heterozygous *MYH7* SNVs in hiPSCs

CRISPR/Cas9 gene-editing and electroporation were used to biallelically edit the endogenous *MYH7* locus in the CLYBL^dCas9-VPR/dCas9-VPR^ WTC11 hiPSC genetic background as above, with adaptions. Briefly, we used the Neon Transfection System with optimized parameters (26) for the 100 μL tip (1200 V, 20 ms, 2 pulses) to transfect 2 × 10^6^ CLYBL^dCas9-VPR/dCas9-VPR^ hiPSCs with 5 μL of each 30 nM synthetic sgRNA (Synthego) targeting *MYH7* intron 21 and 24, 5 μL of 20 μM *Sp*Cas9 2NLS protein (Synthego), 3 μg of pJet-MYH7-eGFP-PGK-PuroR (WT *MYH7*) plasmid DNA, and 540 ng of pJet-MYH7-mTagBFP2-EF1α-BSD (variant *MYH7*) plasmid DNA libraries for each of the five mutagenized amino acid positions in addition to 300 ng of pJet-MYH7-mTagBFP2-EF1α-BSD (*MYH7* p.E848E n.GAG>GAA; referred to as spike-in) plasmid DNA to ensure representation of at least one synonymous SNV. Cells were co-transfected with 2.5 μg pEF1α-BCL-XL plasmid DNA to encourage survival (25).

### CRISPRa on-target editing retrieval (CRaTER)

To obtain a highly enriched population of on-target, biallelically-edited hiPSCs from cells that underwent gene-editing and partial enrichment with antibiotic selection, we developed and applied <u>CR</u>ISPR<u>a</u> On-<u>T</u>arget <u>E</u>diting <u>R</u>etrieval (CRaTER). Briefly, we used the Neon Transfection System with optimized parameters for the 10 μL tip (1400 V, 20 ms, 1 pulse) to transfect 5 × 10^5^ antibiotics selected gene-edited cells with 1 μL of each of three 30 nM synthetic sgRNA (Synthego; see **Table 2**) targeting approximately 150 bp upstream of the *MYH7* transcription start site. A control transfection was performed under the same conditions without any synthetic sgRNAs. One CRaTER transfection was sufficient to retrieve on-target gene-edited cells after gene-editing of individual variants, whereas two CRaTER transfections of library gene-edited cells were combined to maximize retrieval of SNVs. Cells were plated into stem cell media supplemented with CloneR2 and 10 μM ROCK inhibitor without penicillin-streptomycin. Media was replaced the following day and cells allowed to recover for an additional 48 hours.

Cells were harvested 72 hours after transfection by dissociation with 0.5 mM EDTA in PBS, resuspension in stem cell media, filtration using a 70 μm strainer, centrifugation for 3 minutes at 300 x *g*, and resuspension in 200 μL sorting buffer: DPBS, 1% penicillin-streptomycin, 2% FBS, 10 mM HEPES (Invitrogen; cat. no. 15630-080), and 25 μU DNAse I. Cells were stringently sorted into an Eppendorf tube containing stem cell media with 10 μM ROCK inhibitor and CloneR2 using an FACSAria III Cell Sorter (sort speed 1; BD) on the basis of eGFP and mTagBFP2 double-positivity (using the control transfection to set gating). Sorted cells were centrifuged for 3 minutes at 300 × *g* and resuspended and replated in stem cell media with 10 μM ROCK inhibitor and CloneR2. Media was replaced the following day and cells allowed to recover for an additional 96 hours with feeding every other day. Cells were then expanded and a cell pellet was harvested for post-CRaTER bulk genotyping, split at 88 cells/cm^2^ into a 10-cm plate for colony genotyping, and used for cardiac-directed differentiations.

### Genotyping

For single clone isolation, 5,000 hiPSCs were plated in a Matrigel-coated 10-cm plate and cultured for 7 days. Single colonies were manually picked and seeded into a 96-well plate. After 3 days, the hiPSCs were dispersed and seeded into one well of 24-well plate for maintenance and one well of 96-well plate for genotyping. After 3 days, genomic DNA (gDNA) was isolated from hiPSCs with the Quick-DNA Miniprep Kit (Zymo Research; cat. no. D3025) following manufacturer’s instructions. To simultaneously amplify the knock-in and/or un-edited, endogenous allele, PCR genotyping was performed with primers located outside the 5’ homology arm and within *MYH7* exon 23. Clones amplifying only knock-in product without the endogenous allele were further genotyped across separate reactions with a forward primer located outside the 5’ homology arm and a reverse primer specifically binding to either eGFP or mTagBFP2. Clones positive for eGFP and mTagBFP2 were identified and any SNVs were confirmed by Sanger sequencing of the mTagBFP2 PCR product.

### Karyotyping

5 × 10^5^ cells from single variant and variant library hiPSC lines were collected during passaging and genomic DNA was isolated using the DNeasy Blood & Tissue Kit following manufacturer’s instructions (Qiagen; cat. no. 69504). Common karyotypic abnormalities were measured using the qPCR-based hPSC Genetic Analysis Kit following manufacturer’s instructions (STEMCELL Technologies; cat. no. 07550). Cell lines with true karyotypic abnormalities were not used in this study.

### Next generation sequencing (NGS)

#### Plasmid DNA (pDNA) amplicon preparation

DNA amplicons for NGS of each mutagenized plasmid repair template were prepared individually for each position. Briefly, in the first PCR (PCR 1), a 50 μL reaction including 10 ng of pDNA, 10 μM each of common forward and reverse adapter primers (see **Table 1**), and Q5 polymerase were mixed and split into five, 10 μL reactions to minimize jackpotting effects during amplification in a 6 cycle PCR. Sub-reactions were mixed and purification was performed using 30 μL AMPure XP magnetic beads (Beckman Coulter; cat. no. A63880) according to manufacturer’s instructions and eluted in 21 μL H_2_O. In the second PCR (PCR 2), a 50 μL reaction including 2 μL of purified PCR 1 product, 10 μM each of a common forward (F17) and a unique reverse indexing primer (see **Table 1**), and Q5 polymerase were mixed and amplified in a 10 cycle PCR. Purification was performed using 30 μL AMPure XP magnetic beads and eluted in 31 μL H2O. The DNA concentration of individually prepared libraries were quantified using the Qubit dsDNA High Sensitivity Kit (ThermoFisher; cat. no. Q32851) and adjusted to 2 nM before combining for NGS.

#### hiPSC and cardiomyocyte genomic DNA (gDNA) amplicon preparation

Following genomic DNA isolation using the DNeasy Blood & Tissue Kit following manufacturer’s instructions (Qiagen; cat. no. 69504), amplicons were prepared as above, with adaptions only to PCR 1: Briefly, a 50 μL reaction including 50-225 ng of gDNA, 10 μM each of common forward and reverse adapter primers (see **Table 1**), and Q5 polymerase were mixed and split into five, 10 μL reactions for amplification in a 23 cycle PCR.

#### Sequencing parameters

Sequencing was performed on a MiSeq instrument using a MiSeq Reagent Kit v3 (150-cycle; Illumina; cat. no. MS-102-3001) following standard procedures. Custom index and read primers (**Table 1**) were loaded to sequence the amplicons, with 8 cycles allocated for each index and 60 cycles for each read 1 and read 2 sequences. To increase sequence diversity and improve base calling, 10-20% PhiX control DNA (Illumina; cat. no. FC-110-3001) was spiked into the pooled 2 nM DNA amplicon library. To optimize clustering and read output, kits were loaded with a total of 22 pmol of the DNA amplicon library.

#### Data analysis

After demultiplexing on index reads using bcl2fastq version 2.20, paired-end 57-bp reads were merged into a q30 consensus using Paired-End reAd mergeR 0.9.11 and counted (27). pDNA variants with >100 counts (average ~50,000) passed threshold. pDNA variant frequency was calculated as (variant count)/(total variant counts) on a position-by-position basis with only NNK variant codon considered.

NGS of the MYH7^WT-eGFP/WT-mTagBFP2^ hiPSC line (without any SNV) established a threshold for PCR and sequencing error for variants with frequencies less than 0.0004, i.e., hiPSC library variants at a frequency >0.0004 passed threshold. hiPSC variant frequency was calculated as (variant count)/(total variant counts) with only NNK variant codons considered; reads with the WT sequence were removed because our amplicon generation strategy is not allele-specific, therefore we were unable to distinguish between the eGFP and mTagBFP2 alleles.

Cardiomyocyte variants passed threshold if they were present in the hiPSC library (after hiPSC-specific thresholding). To increase stringency, the least frequent 10% of variants were excluded. Cardiomyocyte variant frequency was calculated as (variant count)/(total variant counts) with only NNK variant codons considered.

### Confocal microscopy

Spinning disk microscopy was performed on a Nikon Eclipse Ti equipped with a Yokogawa CSU-W1 spinning disk head, Andor iXon LifeEMCCD camera, and ×100 Plan Apo objective. Images were taken at 37° C and 5% CO_2_.

### Statistical analyses

Bar graphs show mean and error bars indicate standard error mean.

## RESULTS

### CRaTER Enriches for On-Target Gene-Edited hiPSCs

We first examined if CRaTER can be used to enrich for on-target gene-edited hiPSCs by performing HDR in a transcriptionally silent locus (**Figure 1**). To facilitate CRISPRa overexpression during CRaTER, we generated a clonal hiPSC line (WTC11 background) with homozygous knock-in of CAG-dCas9-VPR-P2A-mCherry into the *CLYBL* safe harbor locus (CLYBL^dCas9-VPR/dCas9-VPR^) (**Figure S1**). These cells were then used to biallelically knock-in transgenes containing a promoterless partial *MYH7* cDNA (exon 22-40) fused with a C-terminal fluorescent protein gene (eGFP or mTagBFP2) in *cis* with an antibiotic resistance cassette (PGK-PuroR or EF1α-BSD) into the endogenous *MYH7* locus (**Figure 1B** and **S2A**). These edited cells were allowed to recover and a fraction was collected before and after two days of selection with puromycin and blasticidin for bulk genotyping; a fraction of antibiotics-selected cells was also plated at low density for colony genotyping. Because *MYH7* is a myofilament gene not expressed in pluripotency (15), fluorescence-based selection of knock-in of the transgenic construct is not possible in hiPSCs under normal conditions. Therefore, the remaining edited cells were transfected with three synthetic sgRNA targeting <200 bp upstream the endogenous *MYH7* transcription start site (TSS) in order to transiently drive *MYH7* expression using CRISPRa (a no sgRNA transfection was used as a negative control). Three days after transfection, each condition was harvested for flow cytometric detection of eGFP and mTagBFP2, using the negative control to set gating (**Figure 1C**). eGFP^+^/mTagBFP2^+^ cells were sorted, replated, and allowed to recover and then were harvested for bulk and colony genotyping as before.

**Figure 1.**
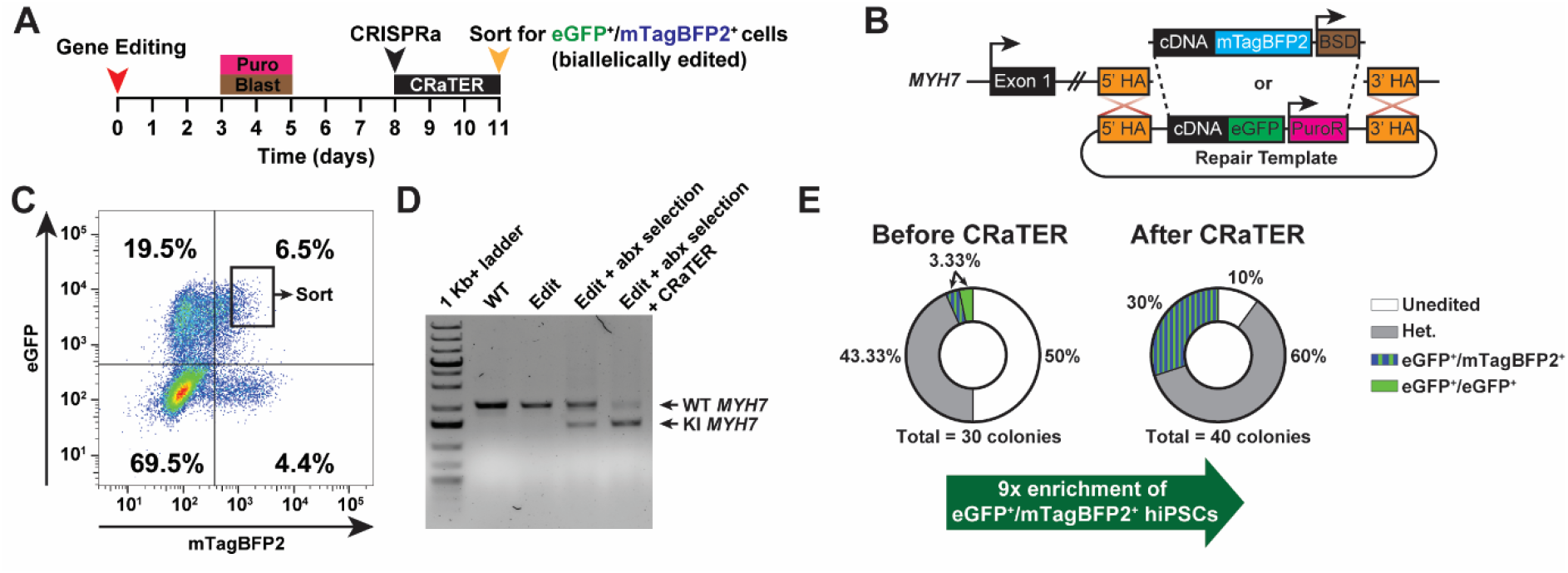
CRaTER enriches for hiPSCs with a biallelically-edited, heterozygous variant knocked into the endogenous *MYH7* locus. (**A**) Schematic of gene-editing and enrichment strategy with <u>CR</u>ISPR<u>a</u> On-<u>T</u>arget <u>E</u>diting <u>R</u>etrieval (CRaTER). Human induced pluripotent stem cells (hiPSCs) stably expressing dCas9-VPR from the *CLYBL* locus (see **Figure S1**) are edited via transfection with Cas9 protein, sgRNA targeting *MYH7* intron 21 and 24, and equal proportions of pJet-MYH7-eGFP-PGK-PuroR (WT *MYH7*) and pJet-MYH7-mTagBFP2-EF1α-BSD (E848E [GAG>GAA] *MYH7*) plasmid DNA repair templates (see **Figure 1B** and **S2A**). Edited cells undergo two days of antibiotic selection and then are transfected with three sgRNA targeting the *MYH7* transcription start site (TSS) to transiently overexpress *MYH7*. Three days later, eGFP^+^/mTagBFP2^+^ hiPSCs (MYH7^WT-eGFP/E848E(GAG>GAA)-mTagBFP2^) are sorted by fluorescence-activated cell sorting (FACS). (**B**) Homology directed repair of the endogenous *MYH7* locus with either plasmid DNA repair template (see **Figure S2**). HA= homology arm; eGFP= enhanced green fluorescent protein; pA= polyadenylation signal; PGK= mouse phosphoglycerate kinase 1 promoter; PuroR= puromycin N-acetyltransferase; mTagBFP2= enhanced monomeric blue fluorescent protein; EF1α= promoter for human elongation factor EF-1α, and BSD= blasticidin S deaminase. (**C**) Flow cytometry plot comparing fluorescence intensities of mTagBFP2 and eGFP in post-antibiotic selected, gene-edited hiPSCs three days after CRISPRa transfection. Sorted eGFP^+^/mTagBFP2^+^ hiPSCs shown in black box. (**D**) Bulk genotyping of unedited WT WTC11 hiPSCs (WT), biallelically-edited CLYBL^dCas9-VPR/dCas9-VPR^ hiPSCs before (Edit) and after (Edit + abx selection) antibiotics selection, and biallelically edited CLYBL^dCas9-VPR/dCas9-VPR^ hiPSCs after antibiotics selection and CRaTER enrichment (Edit + abx selection + CRaTER). The unedited (WT; 2060 bp) and knock-in (KI; 1625 bp) *MYH7* products were specifically amplified from the endogenous *MYH7* locus (see **Figure S2A**). (**E**) Individual colonies from one experiment of post-antibiotic selected, gene-edited hiPSCs were isolated for genotyping the *MYH7* locus before (left) and after (right) CRaTER enrichment. Unedited= biallelic eGFP^+^/mTagBFP2^+^ knock-in without any SNV; het. = one unedited and one knock-in allele.

Bulk PCR-genotyping of the endogenous *MYH7* locus failed to amplify knock-in of the transgenes in the editing only condition, indicating gene-editing was initially rare, however, selection with antibiotics substantially enriched on-target gene-edited cells to 3.3% of the bulk population (**Figure 1D-E**). Moreover, bulk genotyping of eGFP^+^/mTagBFP2^+^ cells sorted after CRaTER enrichment showed preferential amplification of the on-target knock-in *MYH7* transgene relative to endogenous *MYH7*, indicating substantial enrichment for cells with editing of at least one allele (**Figure 1D**). Indeed, genotyping of hiPSCs clones revealed a 9-fold enrichment for eGFP^+^/mTagBFP2^+^ cells after CRaTER enrichment relative to antibiotics selection alone (**Figure 1E**). Together, these data indicate using CRaTER in conjunction with standard antibiotic selection further enriches for rare gene-edited cells.

### CRaTER Enables Efficient Pooled Generation of Over 100 Heterozygous SNVs in hiPSCs

The need to validate on-target gene-editing by isolating and genotyping colonies is a major bottleneck to generating genetic variants in hiPSCs at scale. We hypothesized that we could leverage CRaTER to enrich for on-target heterozygous knock-in of a library of *MYH7* SNVs in hiPSCs and thereby substantially reduce the amount of colony genotyping needed to generate numerous variants. To this end, we targeted five amino acid positions within a hotspot of pathogenic mutations in *MYH7* exon 22 (amino acids 848, 850, 852, 865, and 866) (**Figure 2A**) and mutagenized these positions to saturation in the pJet-MYH7-mTagBFP2-EF1α-BSD plasmid DNA donor repair template. Next generation sequencing (NGS) of these mutagenized plasmids revealed we generated 159/160 (99.4%) possible SNVs at approximately equal frequencies (**Figure S3** and **Table S1**). We transfected the CLYBL^dCas9-VPR/dCas9-VPR^ hiPSC line with equal molarities of two plasmid DNA repair templates: pJet-MYH7-eGFP-PGK-PuroR (WT *MYH7*) and pJet-MYH7-mTagBFP2-EF1α-BSD (variant *MYH7*; comprising equal molarities of each mutagenized position with spike-in of the p.E848E n.GAG>GAA variant) (**Figure 2B**). These edited cells were selected with antibiotics and sampled for colony and bulk genotyping, then enriched with CRaTER as above (**Figure 2C**).

**Figure 2.**
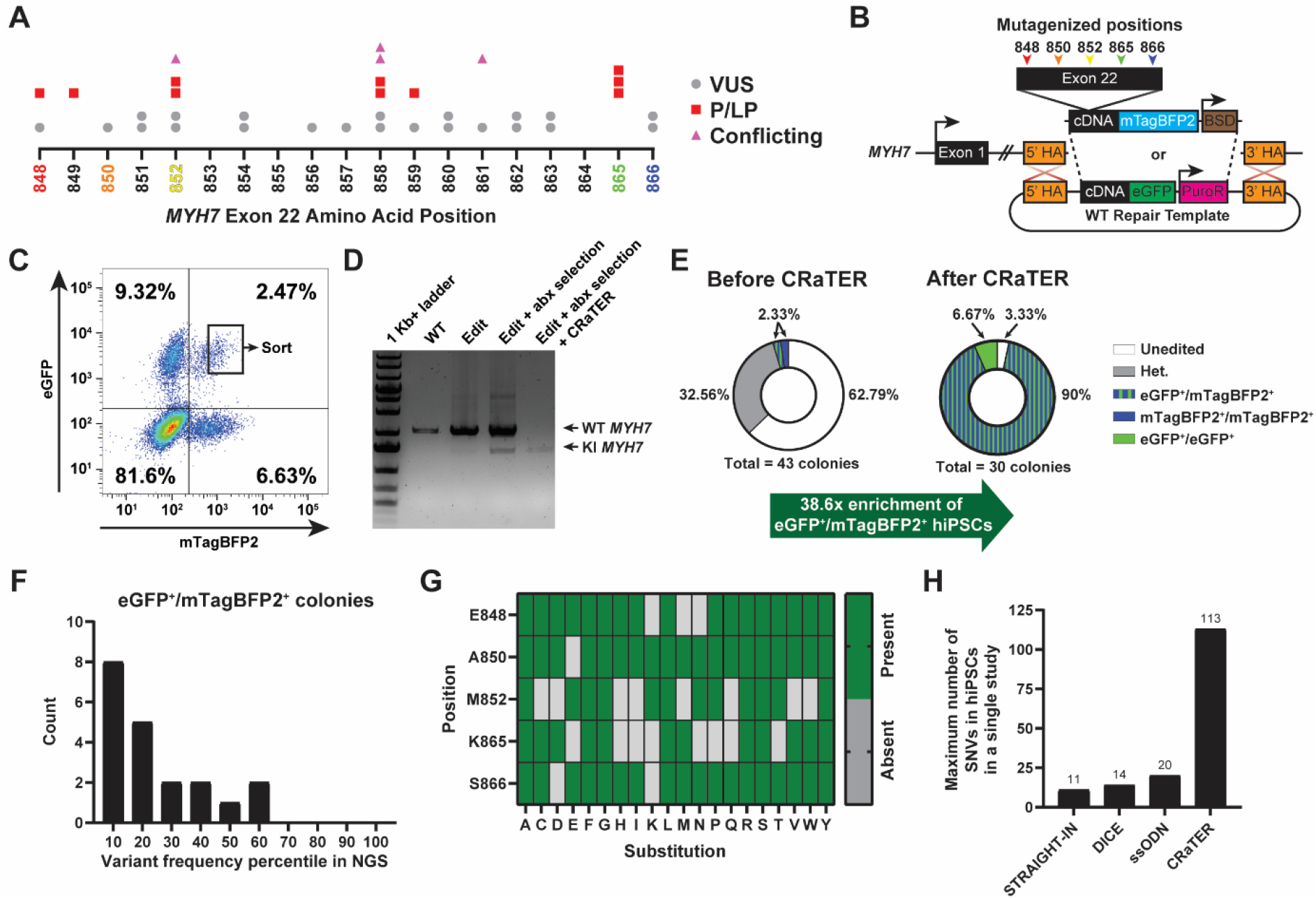
CRaTER enriches for hiPSCs with a biallelically-edited, heterozygous variant library knocked into the endogenous *MYH7* locus. (**A**) Clinical significance of single, heterozygous SNVs within a mutation hotspot region of *MYH7* exon 22 (ClinVar repository as of 1 August 2022). Variants of unknown significance (VUS), pathogenic/likely pathogenic (P/LP), and conflicting interpretations (Conflicting) indicated by grey circles, red squares, and pink triangles, respectively. (**B**) Homology directed repair of the endogenous *MYH7* locus with either plasmid DNA repair template. Within the pJet-MYH7-mTagBFP2-EF1α-BSD repair template only, amino acid positions 848, 850, 852, 865, 866 of *MYH7* exon 22 were mutagenized to saturation. See **Figure S2** for metrics of the pDNA variant library and **Table S1** for associated variant read counts. (**C**) Flow cytometry plot comparing fluorescence intensities of mTagBFP2 and eGFP in post-antibiotic selected, gene-edited hiPSCs three days after CRISPRa transfection. Sorted eGFP^+^/mTagBFP2^+^ hiPSCs shown in black box. (**D**) Bulk genotyping of unedited WT WTC11 hiPSCs (WT), biallelically-edited CLYBL^dCas9-VPR/dCas9-VPR^ hiPSCs before (Edit) and after (Edit + abx selection) antibiotics selection, and biallelically edited CLYBL^dCas9-VPR/dCas9-VPR^ hiPSCs after antibiotics selection and CRaTER enrichment (Edit + abx selection + CRaTER). The unedited (WT; 2060 bp) and knock-in (KI; 1625 bp) *MYH7* products were specifically amplified from the endogenous *MYH7* locus. (**E**) Individual colonies from one experiment of post-antibiotic selected, gene-edited hiPSCs were isolated for genotyping the *MYH7* locus before (left) and after (right) CRaTER enrichment. (**F**) Histogram of isolated and genotyped eGFP^+^/mTagBFP2^+^ hiPSC colonies ranked by frequency in next-generation sequencing (NGS) data and binned by decile. Plot not displaying one variant that failed to pass NGS threshold and six colonies with biallelic knock-in of transgenes without any SNV. See **Figure S3** for metrics of the hiPSC variant library and **Table S2** for associated variant read counts. (**G**) Position map displaying missense variants at targeted positions detected using NGS. Green= present; grey= absent. (**H**) Maximum number of reported SNVs generated in hiPSCs using serine and tyrosine recombinase-assisted integration of genes for high-throughput investigation (STRAIGHT-IN; 11 variants)(40), dual-integrase cassette exchange (DICE; 14 variants) (9), single-stranded oligodeoxyribonucleic (ssODN; 20 variants) (38), and CRaTER (113 variants). Note that the ssODN approach generated homozygous SNVs, whereas DICE and CRaTER (as used here) generates cells with heterozygous SNVs.

Bulk genotyping of the endogenous *MYH7* locus failed to amplify knock-in of the transgenes in the editing only condition, indicating gene-editing was initially rare, however, selection with antibiotics substantially enriched for on-target gene-editing (**Figure 2D**). Moreover, bulk genotyping of eGFP^+^/mTagBFP2^+^ cells sorted after CRaTER enrichment showed a preferential amplification of the on-target knock-in *MYH7* transgene relative to endogenous *MYH7*, indicating substantial enrichment for cells with editing of at least one allele (**Figure 2D**). Indeed, 27/30 (90%) of colonies were eGFP^+^/mTagBFP2^+^ after CRaTER enrichment compared to 1/43 (2.33%) with antibiotics selection alone – a 38.6-fold enrichment (**Figure 2E**). Sanger sequencing of the mutagenized region of *MYH7* in individual eGFP^+^/mTagBFP2^+^ clones revealed 19/27 (70.4%) unique SNVs encoding 18/27 (66.7%) missense variants (**Table 3**). Overall, CRaTER enrichment yielded 18/30 (60%) unique heterozygous *MYH7* missense variants, whereas antibiotics selection alone would require screening ~1,042 colonies (38.6-fold enrichment × 27 eGFP^+^/mTagBFP2^+^ colonies) to yield 18 unique missense variants.

**Table 3.**
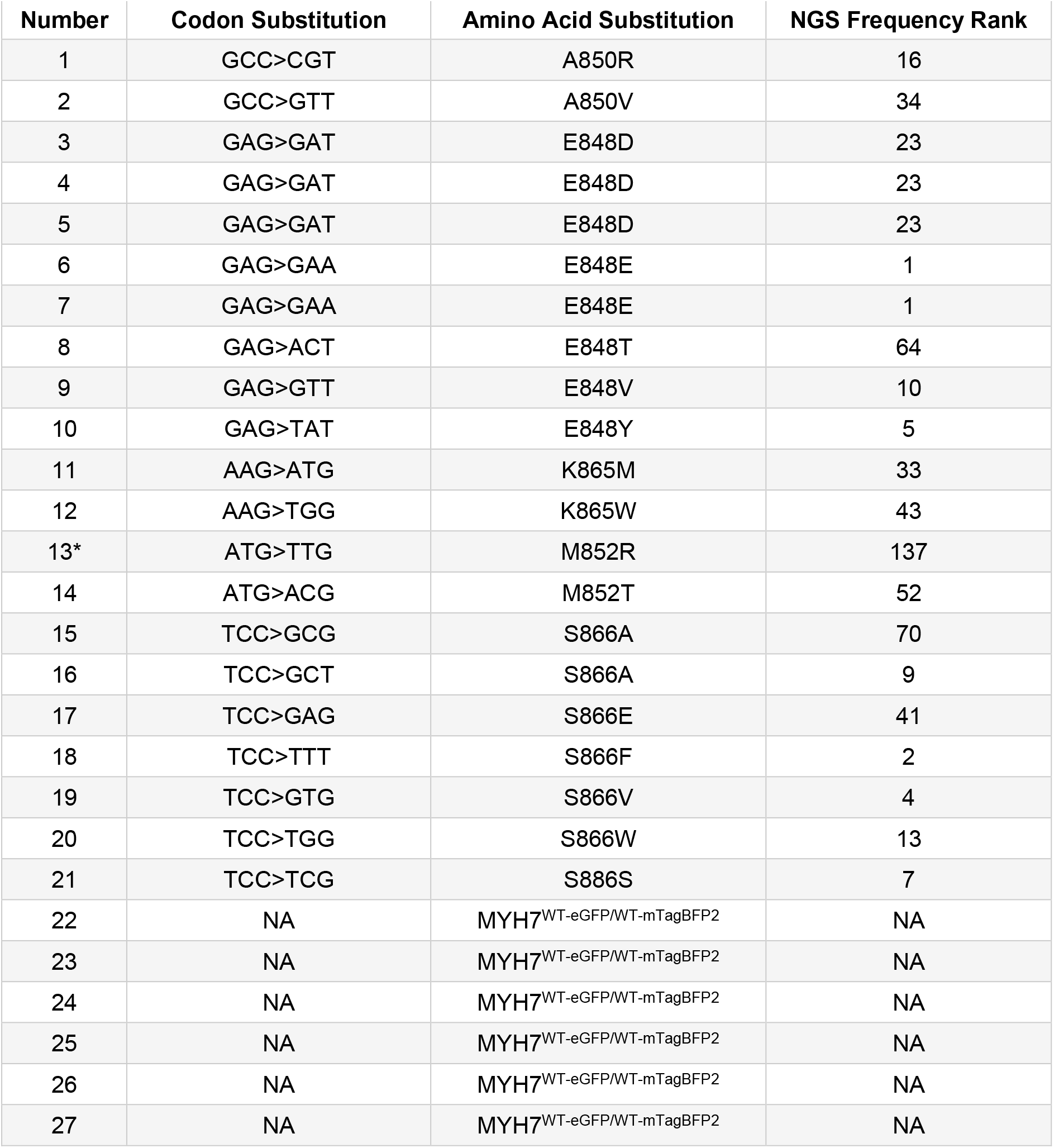
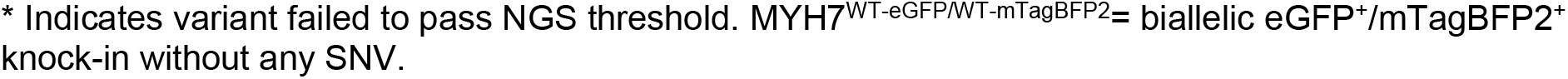
Individual eGFP^+^/mTagBFP2^+^ hiPSC colonies isolated and genotyped after CRaTER.

| Number | Codon Substitution | Amino Acid Substitution | NGS Frequency Rank |
| --- | --- | --- | --- |
| 1 | GCC>CGT | A850R | 16 |
| 2 | GCC>GTT | A850V | 34 |
| 3 | GAG>GAT | E848D | 23 |
| 4 | GAG>GAT | E848D | 23 |
| 5 | GAG>GAT | E848D | 23 |
| 6 | GAG>GAA | E848E | 1 |
| 7 | GAG>GAA | E848E | 1 |
| 8 | GAG>ACT | E848T | 64 |
| 9 | GAG>GTT | E848V | 10 |
| 10 | GAG>TAT | E848Y | 5 |
| 11 | AAG>ATG | K865M | 33 |
| 12 | AAG>TGG | K865W | 43 |
| 13* | ATG>TTG | M852R | 137 |
| 14 | ATG>ACG | M852T | 52 |
| 15 | TCC>GCG | S866A | 70 |
| 16 | TCC>GCT | S866A | 9 |
| 17 | TCC>GAG | S866E | 41 |
| 18 | TCC>TTT | S866F | 2 |
| 19 | TCC>GTG | S866V | 4 |
| 20 | TCC>TGG | S866W | 13 |
| 21 | TCC>TCG | S886S | 7 |
| 22 | NA | MYH7<sup>WT-eGFP/WT-mTagBFP2</sup> | NA |
| 23 | NA | MYH7<sup>WT-eGFP/WT-mTagBFP2</sup> | NA |
| 24 | NA | MYH7<sup>WT-eGFP/WT-mTagBFP2</sup> | NA |
| 25 | NA | MYH7<sup>WT-eGFP/WT-mTagBFP2</sup> | NA |
| 26 | NA | MYH7<sup>WT-eGFP/WT-mTagBFP2</sup> | NA |
| 27 | NA | MYH7<sup>WT-eGFP/WT-mTagBFP2</sup> | NA |

\* Indicates variant failed to pass NGS threshold. MYH7<sup>WT-eGFP/WT-mTagBFP2</sup>= biallelic eGFP<sup>+</sup>/mTagBFP2<sup>+</sup> knock-in without any SNV.

NGS of this mutagenized hiPSC library revealed the generation of 113/157 (72%) SNVs (**Figure S4** and **Table S2**). As expected, the isolated and genotyped hiPSC colonies were among the most frequent variants in the mutagenized hiPSC library (**Figure 2F** and **Table 3**), however, one identified clone did not pass threshold in NGS, suggesting the quality control threshold used in NGS was likely more stringent than necessary. As expected from an overabundance of WT sequences in the mutagenized plasmid library (**Figure S3**), 6/27 (22.22%) of biallelically-edited cells isolated for genotyping were MYH7^WT-eGFP/WT-mTagBFP2^, suggesting incomplete digestion of the plasmid DNA template with DpnI during variant generation (see **Methods** and **Materials**). Overall, we generated 78/97 (80.4%) possible missense variants in heterozygous, biallelically-edited hiPSCs (**Figure 2G**). CRaTER enables the generation of more variants in a single study than all current state-of-the-art gene-editing strategies in hiPSCs (**Figure 2H**). Together, these results indicate CRaTER can be used in conjunction with standard antibiotic selection to further enrich for rare gene-edited cell populations, enabling the pooled generation of variant library hiPSCs at large scale.

### CRaTER Enriched hiPSCs Retain Pluripotency and Differentiate to Functional Cardiomyocytes

We next characterized the differentiation capacity of hiPSCs after CRaTER enrichment and the localization of MHC-β fusion proteins in cardiomyocytes. Using an established, small molecule cardiac-directed differentiation protocol (**Figure 3A**), we differentiated a separately generated MYH7^WT-eGFP/WT-mTagBFP2^ hiPSC line referred to as double knock-in (DKI) from pluripotency to the cardiac lineage. Confocal microscopy of DKI cardiomyocytes 39 days after the onset of differentiation revealed sarcomeric localization of MHC-β-eGFP and MHC-β-mTagBFP2 fusion proteins within the same cell (**Figure 3B**), suggesting these MHC-β fusion proteins do not interfere with normal localization. Moreover, DKI cardiomyocytes spontaneously contract (see **Movie 1**), suggesting basic contractile function of the MHC-β fusion proteins is not impaired.

**Figure 3.**
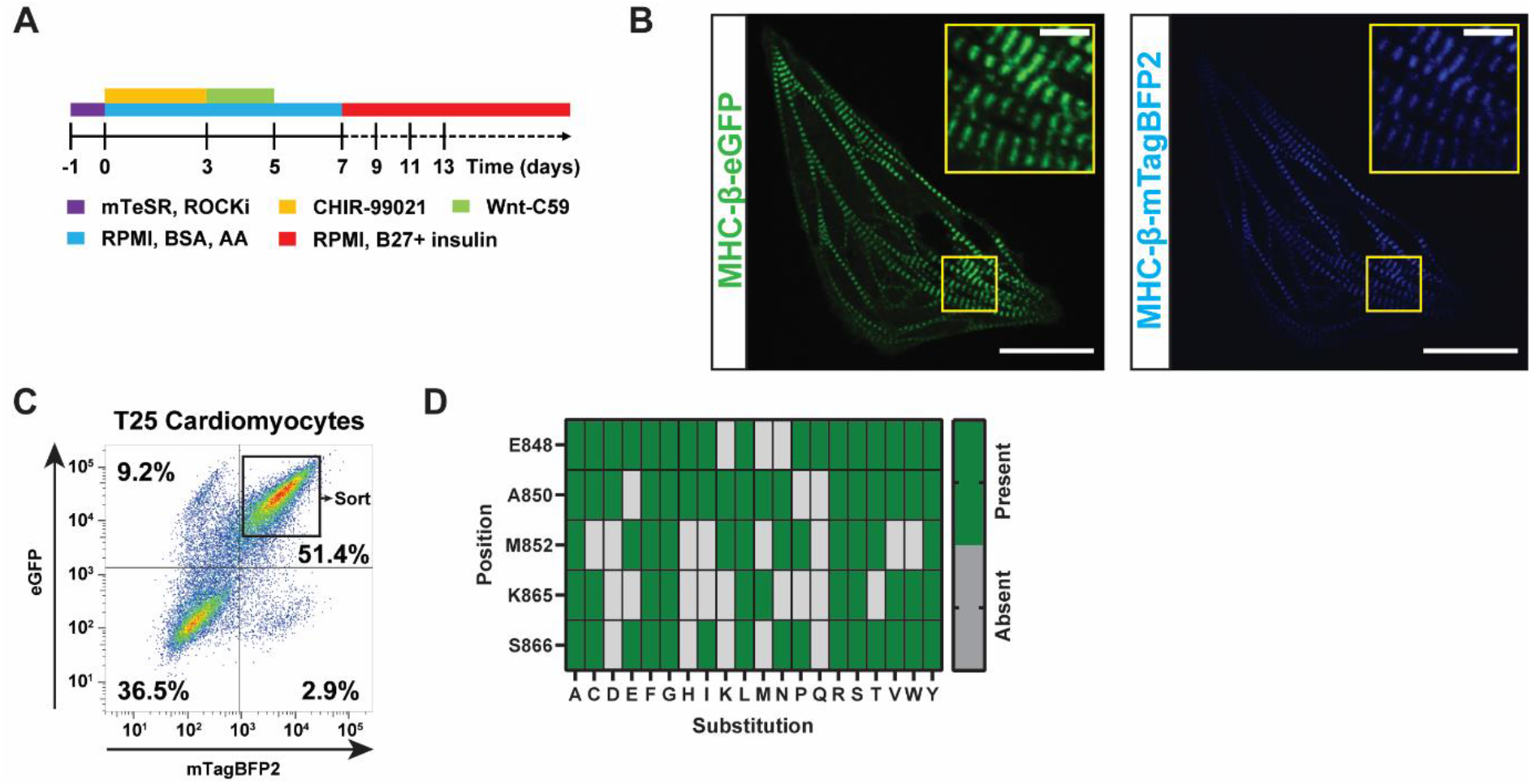
CRaTER enriched hiPSCs differentiate to functional cardiomyocytes and MHC-β fusion proteins localize to sarcomeres. (**A**)Schematic of cardiac-directed differentiation from pluripotency to functional cardiomyocytes. (**B**) Spinning disk confocal micrographs (100x) of live, day 39 cardiomyocytes differentiated from MYH7^WT-eGFP/WT-mTagBFP2^ double knock-in (DKI) hiPSCs. MHC-β-eGFP (green) and MHC-β-mTagBFP2 (blue) expression shown in left and right images, respectively. Main scale bar= 25 μm; inset scale bar= 5 μm. See **Movie S1** for video of spontaneous cardiomyocyte contraction. (**C**) Flow cytometry plot comparing fluorescence intensities of mTagBFP2 and eGFP in day 25 cardiomyocytes differentiated from variant library hiPSCs (from **Figure 2B-F**). Sorted eGFP^+^/mTagBFP2^+^ cardiomyocytes shown in black box. (**D**) Position map displaying missense variants at targeted amino acid positions detected using NGS of day 25 sorted cardiomyocytes (from **Figure 3D**). Green= present; grey= absent. See **Figure S5** for metrics of the variant library and **Table S3** for variant read counts in cardiomyocytes.

We next examined whether the library of heterozygous, biallelically-edited variants in hiPSCs could be differentiated in bulk to cardiomyocytes. Indeed, flow cytometric detection of eGFP and mTagBFP2 from day 25 cardiomyocytes revealed at least 63.5% of cells were cardiomyocytes (expressing at least one fluorescently tagged MHC-β protein), though the true cardiomyocyte purity is likely higher as *MYH7* expression is age-dependent (15,28) (**Figure 3C**). Library edited eGFP^+^/mTagBFP2^+^ cardiomyocytes were sorted and NGS of these mutagenized cardiomyocytes revealed the presence of 102/113 SNVs generated in hiPSCs (90.3%) (**Figure S5** and **Table S3**), translating to 72/78 (80.4%) possible amino acid substitutions in heterozygous, biallelically-edited cardiomyocytes (**Figure 3D**). Together, hiPSCs that have undergone CRaTER enrichment remain pluripotent and can be used in directed-differentiation to generate disease-relevant cell types with minimal variant drop-out following differentiation. In addition, MHC-β-fusion proteins generated using this gene-editing strategy do not cause major defects in MHC-β localization or function in cardiomyocytes.

Overall, CRaTER facilitates on average a 25-fold enrichment of on-target gene-edited hiPSCs from a pool of gene-edited cells. This enrichment greatly reduces the rate-limiting screening required to isolate on-target gene-edited cells, enabling the generation of substantially more gene-edited hiPSCs lines compared to contemporary methods.

## DISCUSSION

In tandem with advanced directed differentiation protocols, human pluripotent stem cells are poised for modelling human genetic diseases to study the effect of variants of unknown significance (VUS) (28,29), elucidate pathological mechanisms (30,31), and test candidate drug therapies (32) in diverse, disease-relevant cell types. The “discovery” of human induced pluripotent stem cells (hiPSC) (33) has allowed for generation of countless hiPSC lines with unique, patient-specific genetic backgrounds, enabling the identification of causative mutations in isogenic, mutation-corrected hiPSC pairs (28,34,35). However, gene editing via homology directed repair (HDR) in stem cells is inefficient (3,36) and has limited the scalable generation of hiPSC tools for various applications, often capping the generation and study of unique cell lines to <3 per publication (37). The rate-limiting step in this process is isolating and genotyping individual hiPSC colonies after gene editing to retrieve rare gene-edited cells. Rather than attempting to directly overcome inefficient gene editing by increasing the HDR rate, we developed a method to enrich for correctly-edited cells called <u>CR</u>ISPR<u>a</u> On-<u>T</u>arget <u>E</u>diting <u>R</u>etrieval (CRaTER). CRaTER enables robust enrichment of on-target gene-edits and can be leveraged to generate a saturated library of gene-edited hiPSCs, creating opportunities for studies of variant effect in disease-relevant cell types at unprecedented scale.

Here, we used CRaTER to enrich for biallelic, heterozygous knock-in of transgenes into the endogenous *MYH7* locus, which is not transcribed in hiPSCs. To facilitate robust CRISPRa activity, we generated a cell line stably expressing dCas9-VPR from the *CLYBL* locus, however, we have also used plasmid-based dCas9-VPR delivery to enrich for WT WTC hiPSCs with on-target knock-in of *MYBPC3*-meGFP (transcriptionally inactive in hiPSCs; data available upon request), indicating CRaTER enrichment is possible in various genetic contexts (*dCas9-VPR* mRNA is also commercially available). We show across independent experiments that CRaTER enables robust enrichment (on average 25-fold higher than antibiotic selection alone) of on-target gene-edited hiPSCs. From a single gene-editing transfection reaction of two million hiPSCs followed by CRaTER enrichment, ~90% of the cells had biallelically-edited heterozygous variants (compared to 2.33% for antibiotic selection alone), consisting of 113 SNVs comprising 78 missense variants. These data suggest the initial biallelic gene-editing efficiency is at least 0.006% (or 113 in 2 million cells), though this is likely an underestimate given that some variants were likely independently knocked in on multiple occasions. Importantly, CRaTER enrichment enables at least a 5-fold greater generation of variants in hiPSCs relative to contemporary state-of-the-art gene-editing approaches (9,38). Together, these results demonstrate that depending on the goal of the end user’s gene-editing, CRaTER can massively reduce the need to isolate and genotype individual colonies. While CRaTER was leveraged here in hiPSCs to exemplify enrichment of rare gene-editing events, CRaTER theoretically could be used to enrich for on-target gene-editing in any cell line amenable to transfection (e.g. HEK 293T, HAP1, and U-2 OS).

CRaTER is designed to be applied for gene-editing enrichment when the targeted gene of interest (GOI) is not expressed, or is lowly expressed. CRISPRa targeting of the endogenous GOI locus in gene-edited cells transiently and specifically overexpresses the knocked-in transgene, allowing for fluorescent detection of cells with correctly-edited genes and sorting using FACS. If a similar gene-editing strategy is pursued in a GOI that is already robustly expressed in hiPSCs, then CRaTER is likely not necessary because the on-target gene-edited cells will express the fluorescent protein and can be directly sorted. Transient overexpression of certain GOIs, such as transcription factors, during CRISPRa could potentially initiate differentiation in hiPSCs and should be carefully considered to avoid loss of pluripotency. Additionally, overexpression of certain GOIs could be cytotoxic when aberrantly expressed.

While CRISPR/Cas9 editing was used here, theoretically any HDR-based gene-editing strategy could be used. We intentionally knocked-in a partial cDNA-fluorescent protein fusion transgene to demonstrate that the MHC-β-fluorescent protein localizes and functions normally in cardiomyocytes, however, CRaTER theoretically could be used to enrich for various gene-editing strategies using different transgenes, such as: (1) a self-cleaving partial cDNA-T2A-fluorescent protein gene (if the fusion affects normal biology), (2) a partial cDNA-T2A-antibiotics resistance gene (if the cell type does not tolerate sorting), or (3) a C-terminal fluorescent protein gene alone (if a simple reporter is desired). Note that knock-in of complete cDNA fused with a C-terminal fluorescent protein gene could initiate HUSH silencing due to lack of introns (39), suggesting that the gene-editing strategy used with CRaTER should target downstream of the initial exons to minimize HUSH activity.

We show that manual deconvolution of the CRaTER enriched variant hiPSC library is a feasible strategy to retrieve numerous unique missense variants for individual characterization. CRaTER enables future studies leveraging high-throughput characterization of pooled cellular libraries to determine variant effect in disease-relevant stem cell-derived cell types.

Together, we report an enrichment method using existing technology in a novel manner to facilitate the generation of transgenic hiPSCs lines by minimizing individual colony genotyping. We expect CRaTER will streamline gene editing in hiPSCs and other cell types for applications in disease modelling.

## Supporting information

Graphical Abstract

## DATA AVAILABILITY and ACCESSION NUMBERS

The data presented in this publication have been deposited in NCBI’s Gene Expression Omnibus (GEO). It is currently private but is available through the GEO Series accession number GSE213520 and the following URL link: https://www.ncbi.nlm.nih.gov/geo/query/acc.cgi?acc=GSE213520

The following reviewer token will allow anonymous, read-only access to GSE213520 while it is private: ylgtaciujfozfyr

GSE213520 will be made publicly-available prior to publication of the manuscript.

## ACKNOWLEDGEMENTS

The WT WTC11 hiPSCs were a gift from Bruce Conklin. The pEF1α-BCL-XL vector was a gift from Xiao-Bing Zhang. The pC13N-iCAG.copGFP, pZT-C13-L1, and pZT-C13-R1 vectors were a gift from Jizhong Zou. The lenti-EF1α-dCas9-VPR-Puro vector was a gift from Kristen Brennand. The IGI-P0492 pHR-dCas9-NLS-VPR-mCherry vector was a gift from Jacob Corn. We are thankful to the staff of the Cell Analysis Facility at the University of Washington.

## AUTHOR CONTRIBUTIONS

Conceptualization, C.E.F., S.F., S.P., L.M.S., D.M.F., and K-C.Y.; Methodology, C.E.F., S.F., W-M.C., and K-C.Y.; Software, S.P.; Validation, C.E.F. and S.P.; Formal Analysis, C.E.F.; Investigation, C.E.F., S.F., S.P., W-M.C., L.T., S-L.C., A.K., and A.M.K; Resources, E.K., L.M.S., and D.M.F.; Data Curation, C.E.F., W-M.C., and S.P.; Writing – Original Draft, C.E.F.; Writing – Review & Editing, C.E.F., S.F., S.P., W-M.C., E.K., A.L. C.E.M., L.M.S., D.M.S., and K-C.Y.; Visualization, C.E.F.; Supervision, C.E.F. and K-C.Y.; Project Administration, K-C.Y.; Funding Acquisition, C.E.F., S.F., S.P., E.K., A.M.F., A.L., C.E.M., L.M.S., D.M.F., and K-C.Y.

## SUPPLEMENTARY DATA

**Figure S1. CLYBL^dCas9-VPR/dCas9-VPR^ transgenesis in WTC11 hiPSCs.**

**Figure S2. Gene-editing of endogenous *MYH7* locus and possible outcomes. Figure S3. Plasmid DNA repair template variant library NGS metrics.**

**Figure S4. hiPSC variant library NGS metrics.**

**Figure S5. Cardiomyocyte variant library NGS metrics.**

**Table S1. Plasmid DNA variant library NGS data.**

**Table S2. hiPSC variant library NGS data.**

**Table S3. Cardiomyocyte variant library NGS data.**

**Movie S1**. Spontaneous day 39 MYH7^WT-eGFP/WT-mTagBFP2^ cardiomyocyte contraction captured with spinning disk confocal microscopy (100X objective). Note photobleaching causes dimming over time.

