## Supplemental Figures for "CRaTER enrichment for on-target gene-editing enables generation of variant libraries in hiPSCs"

### SUPPLEMENTARY DATA

**Figure S1. CLYBL$^{dCas9\text{-}VPR/dCas9\text{-}VPR}$ transgenesis in WTC11 hiPSCs.**
**Figure S2. Gene-editing of endogenous *MYH7* locus and possible outcomes.**
**Figure S3. Plasmid DNA repair template variant library NGS metrics.**
**Figure S4. hiPSC variant library NGS metrics.**
**Figure S5. Cardiomyocyte variant library NGS metrics.**

**Table S1. Plasmid DNA variant library NGS data.**
**Table S2. hiPSC variant library NGS data.**
**Table S3. Cardiomyocyte variant library NGS data.**

**Movie S1.** Spontaneous day 39 MYH7$^{WT\text{-}eGFP/WT\text{-}mTagBFP2}$ cardiomyocyte contraction captured with spinning disk confocal microscopy (100X objective). Note photobleaching causes dimming over time.

Supplementary Data are available at NAR online.

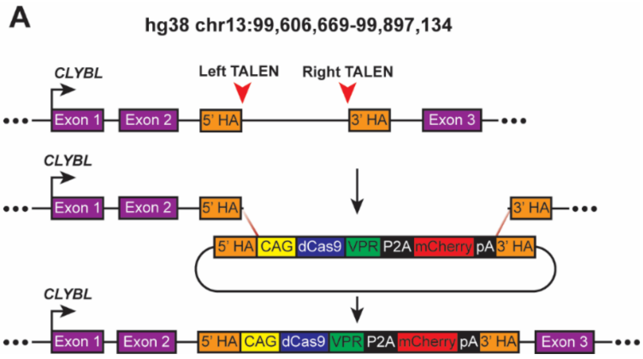


**Figure S1. CLYBL$^{dCas9\text{-}VPR/dCas9\text{-}VPR}$ transgenesis in WTC11 hiPSCs.**
(**A**) Transgenesis strategy of the CAG-dCas9-VPR-P2A-mCherry plasmid DNA repair template construct into the endogenous *CLYBL* locus in hiPSCs. Red arrowheads indicate TALENs targeting between exon 2 and 3. HA= homology arm; CAG= chicken β-actin promoter; dCas9= catalytically dead mutant of the Cas9 endonuclease from the *Streptococcus pyogenes* Type II CRISPR/Cas system; VPR= tetrameric repeat of the minimal activation domain of herpes simplex virus VP16 with transcriptional activation domain of human RelA and Rta transactivator from the Epstein-Barr virus; P2A= 2A peptide from porcine teschovirus-1 polyprotein; mCherry= monomeric derivative of DsRed fluorescent protein, and pA= polyadenylation signal.

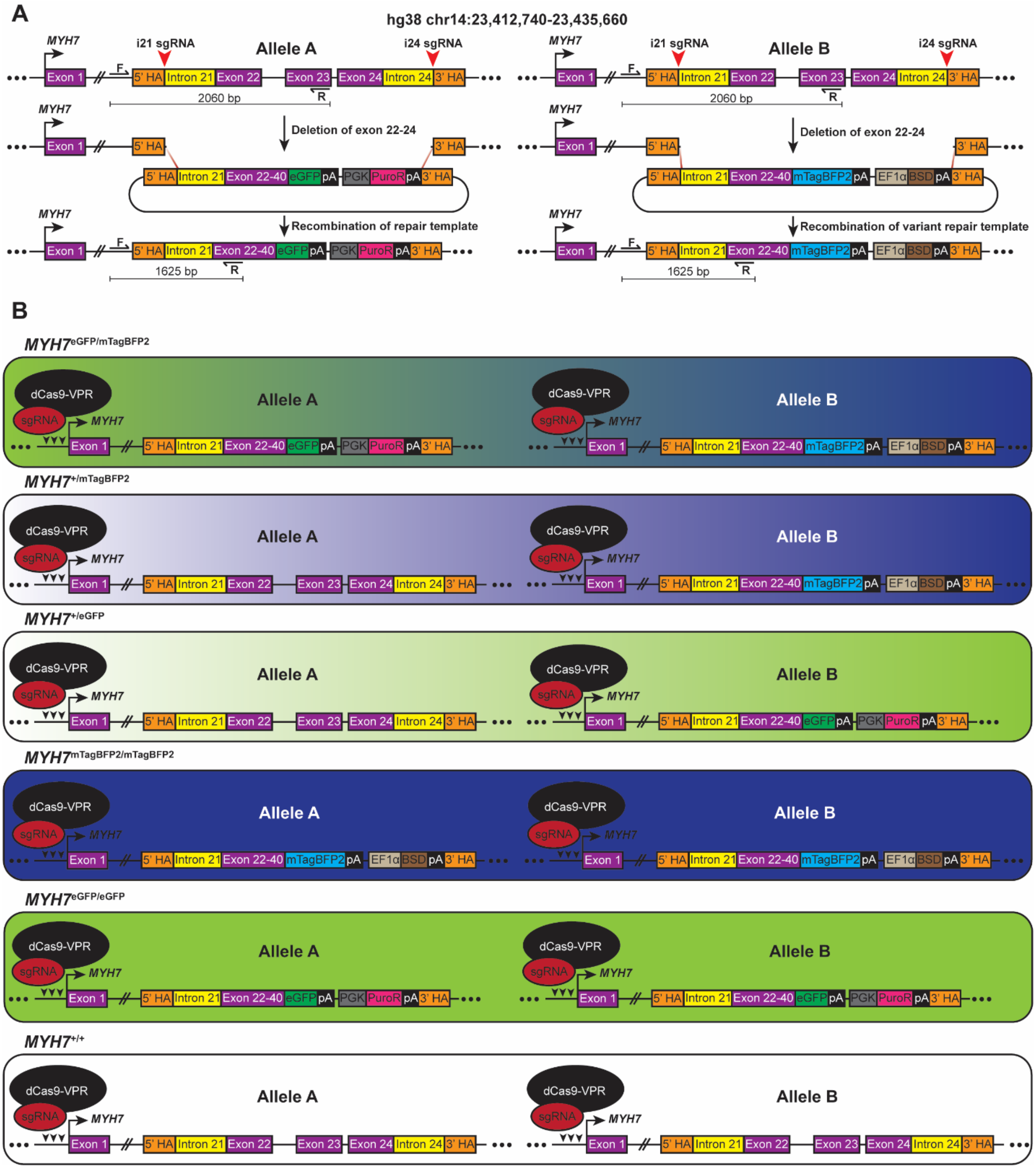


**Figure S2. Gene-editing of endogenous *MYH7* locus and possible outcomes.**
(**A**) Transgenesis strategy of the plasmid DNA repair template constructs into endogenous *MYH7* alleles in hiPSCs. Red arrowheads indicate sgRNA targeting *MYH7* intron 21 and 24. Forward (F) and reverse (R) primers amplify a 2060 bp product with no insertion or a 1625 bp product if the transgene is knocked into the endogenous *MYH7* locus (used for genotyping in **Figure 1D** and **2D**). HA= homology arm; eGFP= enhanced green fluorescent protein; pA= polyadenylation signal; PGK= mouse phosphoglycerate kinase 1 promoter; PuroR= puromycin N-acetyltransferase; mTagBFP2= enhanced monomeric blue fluorescent protein; EF1α= promoter for human elongation factor EF-1α; and BSD= blasticidin S deaminase.
(**B**) Possible outcomes of gene-editing in (**A**) during CRISPRa targeting of the *MYH7* transcription start site.

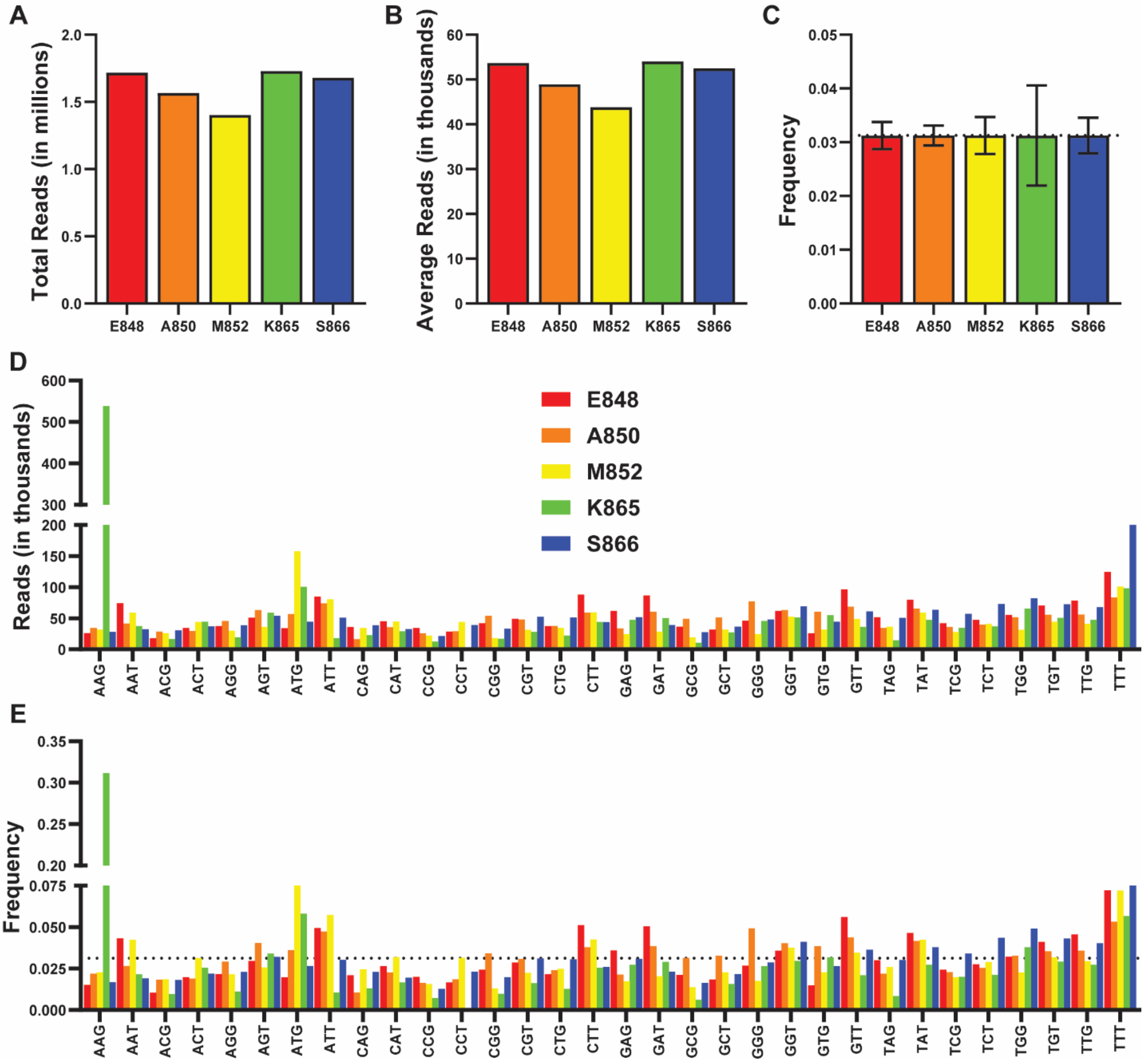


**Figure S3. Plasmid DNA repair template variant library NGS metrics.**
(**A**) Total sequenced reads in millions for mutagenized amino acid positions in *MYH7*. Only data from NNK codons shown in **Figure S3**.
(**B**) Average sequenced reads in thousands for mutagenized amino acid positions in *MYH7*.
(**C**) Frequency of sequenced variants for mutagenized amino acid positions *MYH7*. Dotted line at y= 0.03125 indicates expected 1/32 variant frequency per amino acid position. Error bars indicate standard error mean.
(**D**) Sequenced reads in thousands for each NNK variant within mutagenized amino acid positions in *MYH7*. E848= red; A850= orange; M852= yellow; K865= green; S866= blue.
(**E**) Frequency of sequenced variants for each NNK variant within mutagenized amino acid positions in *MYH7*. Dotted line at y= 0.03125 indicates expected 1/32 variant frequency per amino acid position.

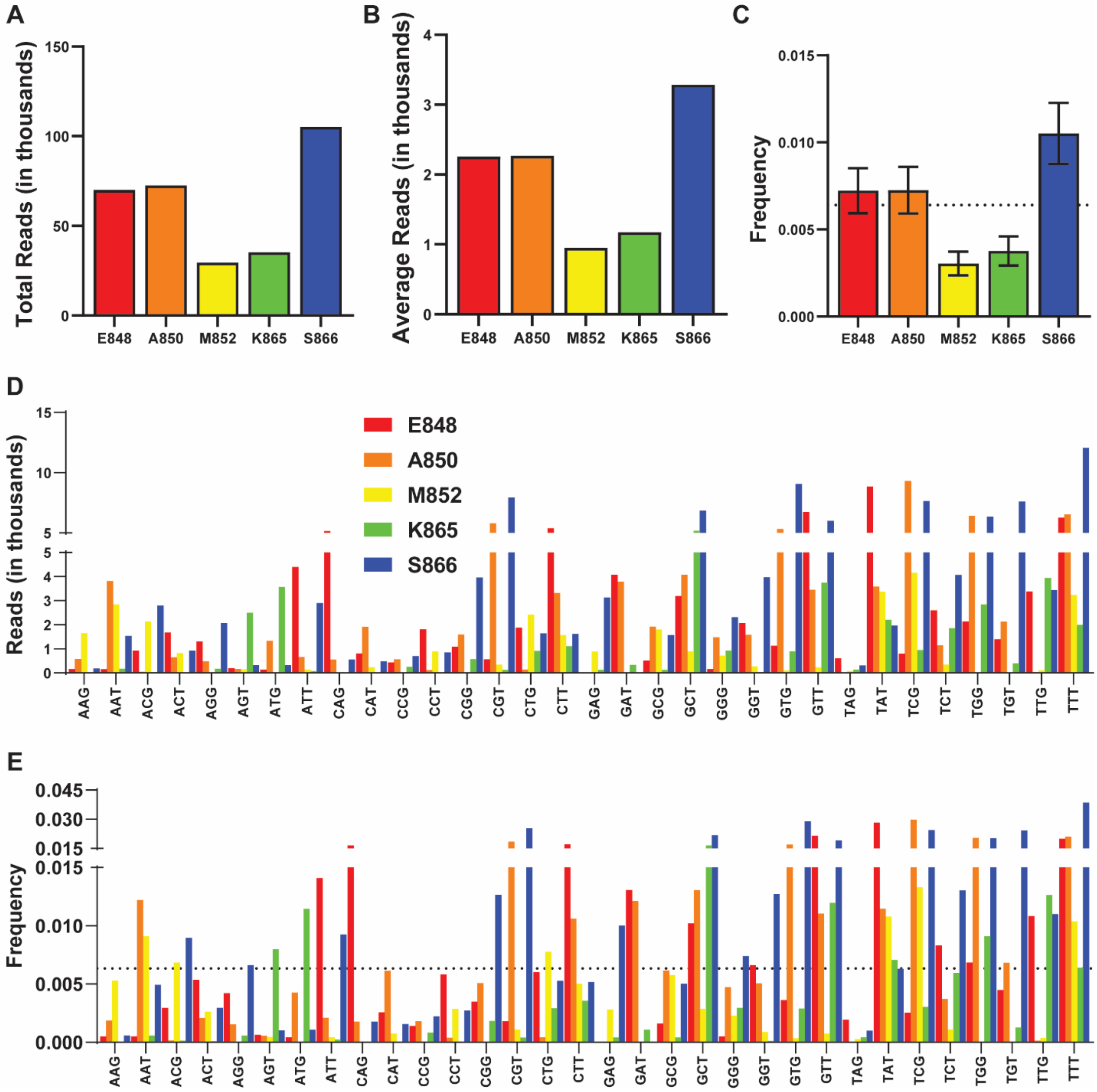


**Figure S4. hiPSC variant library NGS metrics.**

(**A**) Total sequenced reads in thousands for mutagenized amino acid positions in *MYH7*. Only data from NNK codons (wildtype and p.E848E n.GAG>GAA spike-in sequences removed) shown in **Figure S4**.

(**B**) Average sequenced reads in thousands for mutagenized amino acid positions in *MYH7*.

(**C**) Frequency of sequenced variants for mutagenized amino acid positions *MYH7*. Dotted line at y= 0.0064102 indicates expected 1/156 possible variant frequency in library. Error bars indicate standard error mean.

(**D**) Sequenced reads in thousands for each NNK variant within mutagenized amino acid positions in *MYH7*. E848= red; A850= orange; M852= yellow; K865= green; S866= blue.

(**E**) Frequency of sequenced variants for each NNK variant within mutagenized amino acid positions in *MYH7*. Dotted line at y= 0.0064102 indicates expected 1/156 possible variant frequency in library.

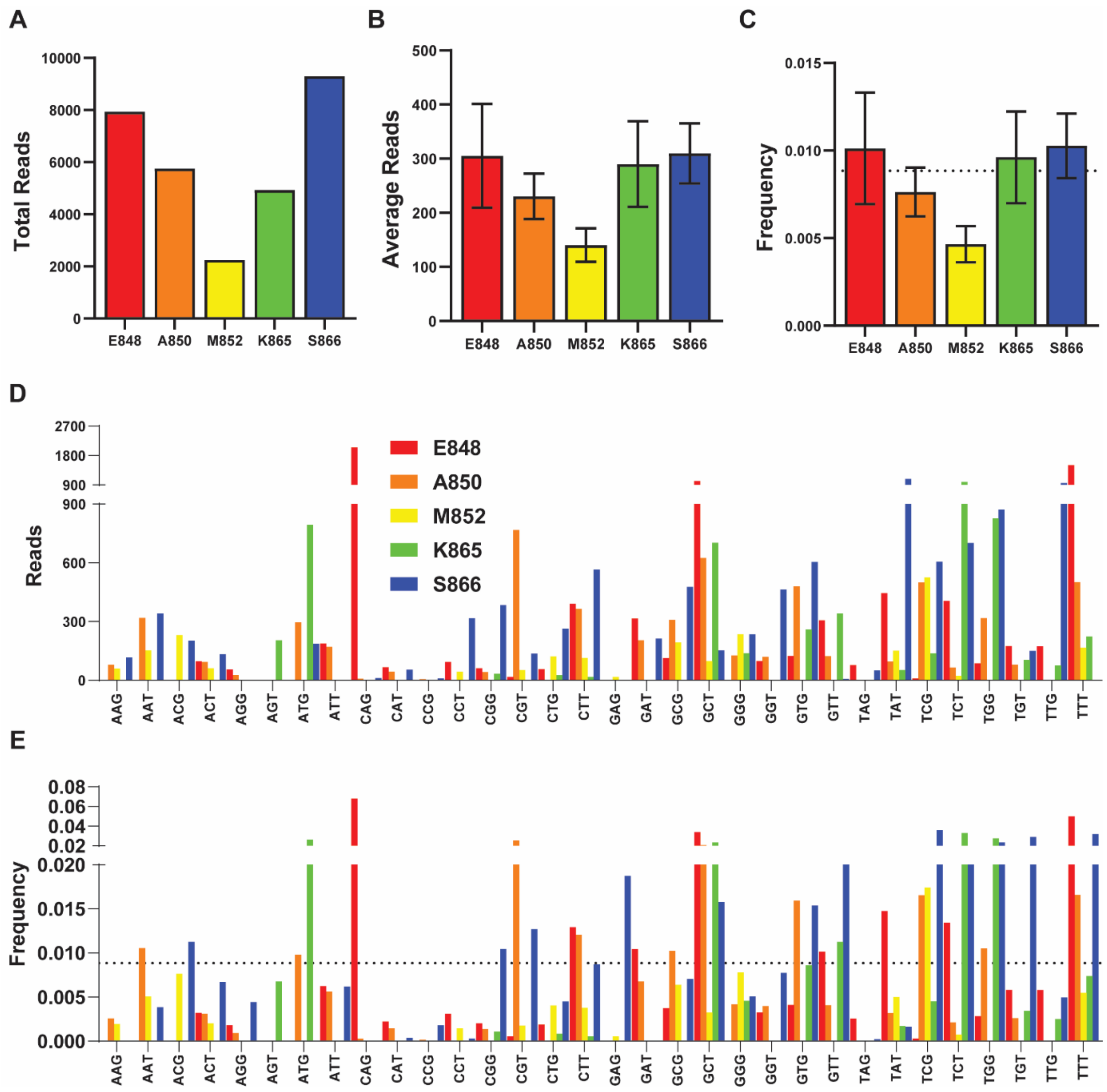


**Figure S5. Cardiomyocyte variant library NGS metrics.**
(**A**) Total sequenced reads in thousands for mutagenized amino acid positions in *MYH7*. Only data from NNK codons (wildtype and p.E848E n.GAG>GAA spike-in sequences removed) shown in **Figure S5**.
(**B**) Average sequenced reads in thousands for mutagenized amino acid positions in *MYH7*.
(**C**) Frequency of sequenced variants for mutagenized amino acid positions *MYH7*. Dotted line at y= 0.008849 indicates expected 1/113 possible variant frequency in library. Error bars indicate standard error mean.
(**D**) Sequenced reads in thousands for each NNK variant within mutagenized amino acid positions in *MYH7*. E848= red; A850= orange; M852= yellow; K865= green; S866= blue.
(**E**) Frequency of sequenced variants for each NNK variant within mutagenized amino acid positions in *MYH7*. Dotted line at y= 0.008849 indicates expected 1/113 possible variant frequency in library.
