## Supplementary material for "CRaTER enrichment for on-target gene-editing enables generation of variant libraries in hiPSCs": Graphical Abstract

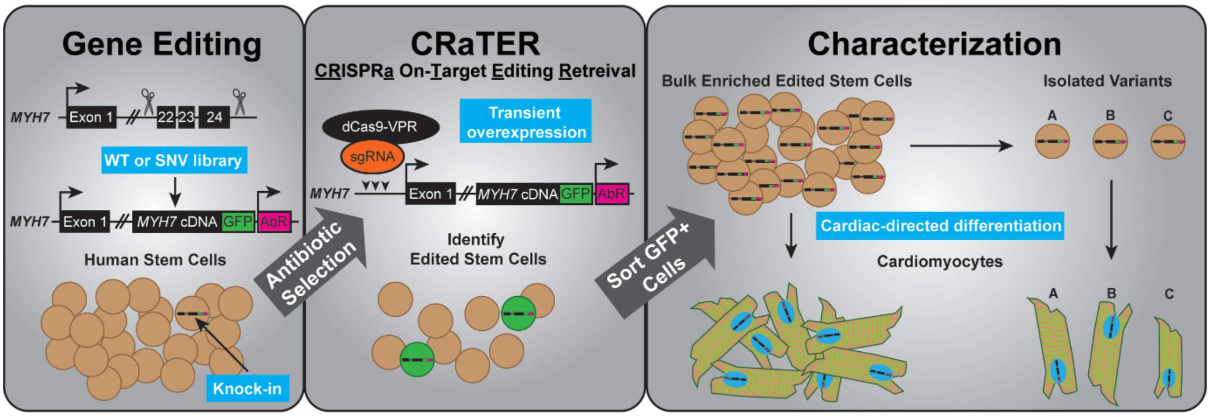


**Graphical Abstract.** On-target gene-edited iPSCs (**left**) are enriched using CRaTER (**middle**), enabling rapid, pooled generation of variant libraries for characterization in relevant cell types of interest after directed differentiation (**right**).
